# Kidney Tissue Characterization using Normalized Raman Imaging and Segment-Anything

**DOI:** 10.1101/2025.10.09.681509

**Authors:** Ranit Karmakar, William V. Trim, Marc Kirschner, Simon F. Nørrelykke

## Abstract

Normalized Raman Imaging (NoRI) enables high-resolution, label-free quantification of protein and lipid concentrations in biological tissues. Because NoRI provides rich molecular information, the analysis of its large, multi-channel datasets turns into a significant computational bottleneck. In this work, we introduce a novel, modular computational pipeline for automated segmentation and quantification of kidney tissue structures imaged with NoRI. The pipeline integrates classical image processing with state-of-the-art machine learning tools, including the Segment Anything Model (SAM) and ilastik, to segment key anatomical and biochemical features—such as tubules, nuclei, brush borders, and lumens. A custom contrast-enhancement strategy was developed to create a third SAM input channel from NoRI data, leading to a substantial improvement in segmentation performance (F1 score: 0.9226). Our framework enables accurate cytoplasm, resolved quantification of protein and lipid concentrations and reveals distinct biochemical signatures across renal tubule subtypes and experimental conditions. This method offers a robust, scalable foundation for quantitative tissue analysis and enhances the utility of NoRI imaging for biomedical research.

## 1. Introduction

Biological tissues exhibit complex spatial variation in protein and lipid composition, which plays a critical role in organ function, development, and pathology. Accurately quantifying and mapping these biochemical properties at subcellular resolution is essential for understanding normal physiology and disease mechanisms across a wide range of tissue types.

Stimulated Raman Scattering (SRS) microscopy has emerged as a powerful, label-free technique for imaging protein and lipid content with high spatial resolution. Normalized Raman Imaging (NoRI), a recent extension of SRS, corrects for signal attenuation in thick samples, enabling absolute quantification of protein and lipid concentrations across entire tissue sections (1). However, its high-resolution, multi-channel datasets are complex, large, and difficult to analyze using traditional tools. Existing segmentation methods do not generalize well to NoRI data, and training new models is impractical—manual annotation is time-consuming and requires specialized domain expertise, making large-scale labeled datasets difficult to obtain.

To address this gap, we present a modular computational pipeline for the automated segmentation and quantification of tissue structures in NoRI images. Rather than designing new deep learning models, our goal is to build a robust and adaptable framework that makes analyzing NoRI data more accessible and useful to researchers. The pipeline combines domain-specific pre-processing to adapt it with state-of-the-art vision tools—including the Segment Anything Model (SAM) (2) and *ilastik*—to enable accurate segmentation with minimal manual input. Importantly, it is designed to run efficiently on standard computing hardware, supporting interactive and scalable analysis.

**The pipeline was developed around four key design goals:**

1. **Modular:** Components can be independently modified or replaced for different segmentation, classification, or quantification tasks.
2. **Efficient:** Runs quickly on consumer-grade GPUs and remains usable on CPUs.
3. **Robust:** Avoids reliance on immunofluorescence (IF) markers; operates solely on NoRI channels.
4. **Generalizable:** Adaptable to new tissue types and experimental conditions with minimal configuration.

We demonstrate the utility of this pipeline on kidney tissue, which presents a diverse set of structural and biochemical features. However, the design and methods are not kidney-specific. **This framework is intended as a general tool for accelerating NoRI-based discovery in a broad range of biological settings**. By lowering the barrier to quantitative analysis, we aim to support wider adoption of NoRI imaging and its application to biological, clinical, and translational research.

## 2. Dataset and Experimental Setup

The dataset used in this study consists of 100 images, with each image having between 6-16 field of views (FoVs) categorized into three experimental groups: male and female kidneys, ischemic and control kidneys, and regional kidney analysis. Each group was designed to address specific biological questions related to kidney function and pathology.

For all images, three channels were consistently captured: two Raman channels representing protein and lipid concentrations, and a third immunofluorescence (IF) channel for endomucin. Additionally, specific markers were applied depending on the experimental group. In the male and female kidney experiments, as well as those focused on regional kidney analysis, three additional markers were used: L-Tumor Lectin (LTL) for proximal tubules, uromodulin for thick ascending limbs, and Aquaporin 2 (AQP2) for collecting ducts. The Raman channels were captured at 16-bit depth, while the IF channels were recorded at 12-bit depth.

For the ischemic kidney experiments, the LTL marker was replaced with a kidney-damage marker Kidney Injury Molecule-1 (KIM1) to facilitate targeted analysis of ischemic injury and its effects on kidney structure and function.

The images varied in (*x, y*) dimension based on the sample size and region of the kidney. Table **1** provides a summary of the number of images per experiment. A detailed description of the dataset and acquisition protocol can be found in Trim et. al. (3).

**Table 1:** Experimental Dataset Overview: Summary of the number of images, fields of view (FoVs) per image, and image dimensions used in the study.

| Group | Number of Images | FoVs/Image | Image Dimensions |
| --- | --- | --- | --- |
| **Kidney Ischemia Timecourse** |  |  |  |
| Control, Day 1 post-ischemia | 10 | 4–6 | 1970 × 1970 px<br>1970 × 2942 px |
| Control, Day 2 post-ischemia | 6 | 4–6 | 1970 × 1970 px<br>1970 × 2942 px |
| Control, Day 5 post-ischemia | 8 | 4–6 | 1970 × 1970 px<br>1970 × 2942 px |
| Control, Day 25 post-ischemia | 7 | 4–6 | 1970 × 1970 px<br>1970 × 2942 px |
| Control, Day 25 post-ischemia (KIM1 stained) | 7 | 4–8 | 1970 × 1970 px<br>1970 × 3914 px |
| Ischemic, Day 1 post-ischemia | 6 | 6–8 | 1970 × 2456 px<br>1970 × 2942 px<br>1970 × 3428 px |
| Ischemic, Day 2 post-ischemia | 7 | 4–6 | 1970 × 1970 px<br>1970 × 2456 px<br>1970 × 2942 px |
| Ischemic, Day 5 post-ischemia | 4 | 4–6 | 1970 × 1970 px<br>1970 × 2942 px |
| Ischemic, Day 25 post-ischemia | 8 | 4–6 | 1970 × 1970 px<br>1970 × 2942 px |
| Ischemic, Day 25 post-ischemia (KIM1 stained) | 8 | 4–8 | 1970 × 1970 px<br>1970 × 3914 px |
| **Sex Differences of Healthy Kidneys** |  |  |  |
| Healthy Kidney (Female) | 5 | 10 | 1970 × 4400 px<br>1970 × 4886 px |
| Healthy Kidney (Male) | 6 | 6–12 | 1970 × 2942 px<br>1970 × 5372 px<br>1970 × 5858 px |
| **Regional Analysis of Healthy Kidneys** |  |  |  |
| Healthy Kidney, 0–1 mm | 2 | 16 | 1970 × 7820 px |
| Healthy Kidney, 1–2 mm | 2 | 16 | 1970 × 7820 px |
| Healthy Kidney, 2–3 mm | 2 | 16 | 1970 × 7820 px |
| Healthy Kidney, 3–4 mm | 2 | 16 | 1970 × 7820 px |
| Healthy Kidney, 4–5 mm | 2 | 16 | 1970 × 7820 px |
| Healthy Kidney, 5–6 mm | 2 | 16 | 1970 × 7820 px |
| Healthy Kidney, 6–7 mm | 2 | 16 | 1970 × 7820 px |
| Healthy Kidney, 7–8 mm | 2 | 16 | 1970 × 7820 px |
| Healthy Kidney, 8–9 mm | 2 | 16 | 1970 × 7820 px |
| **Total** | 100 | – | – |

### 2.1. Ground Truth Creation

Manually generating ground truth segmentations for tubules in NoRI images is a labor-intensive and error-prone process, subject to human variability and inconsistency. To address these challenges and ensure high-quality and reproducible annotations, we implemented a semi-automated workflow that combines machine learning-based segmentation with expert oversight, as illustrated in Figure **1**.

**Fig. 1:**
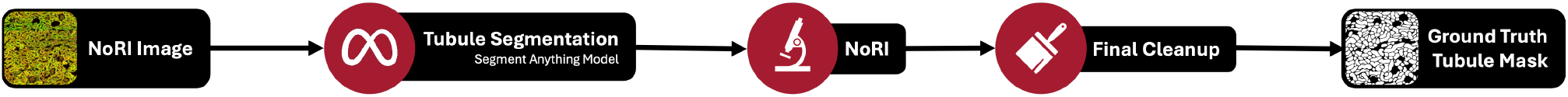
Workflow for generating ground truth tubule segmentations from NoRI images: Ground truth labels were created through a multi-step process. First, the original NoRI images were processed using the Segment Anything Model to obtain initial tubule segmentations. These segmentations were subsequently reviewed and refined by an expert to correct inaccuracies. Finally, a secondary correction step was performed to ensure high-quality, reliable annotations for downstream analysis and model training.

**Fig. 2:**
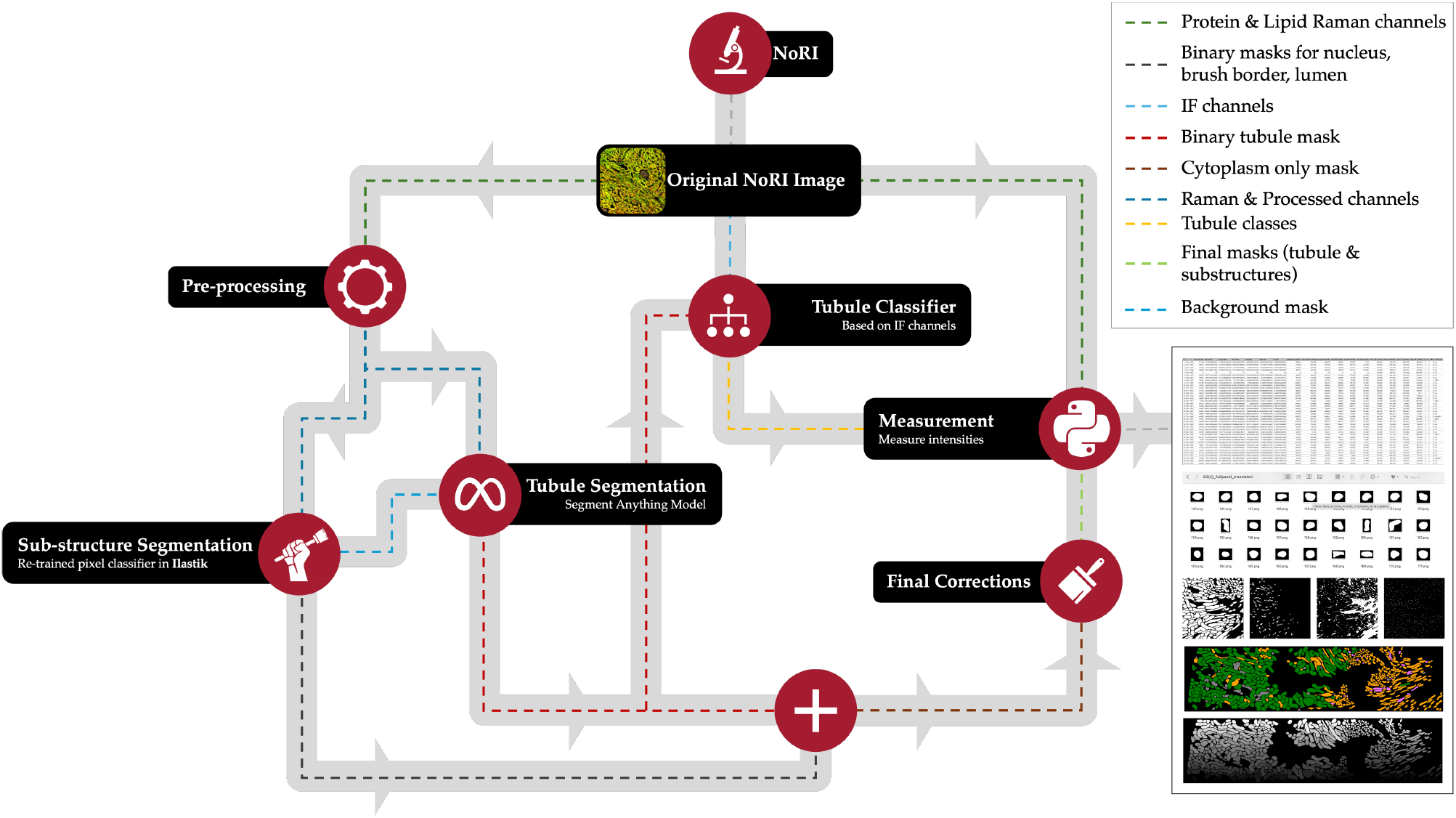
Computational pipeline for segmentation and quantification of protein and lipid concentrations in kidney tissues. Original NoRI images are pre-processed for contrast enhancement. Tubule segmentation is performed using the Segment Anything Model (SAM), and sub-structures (nuclei, brush borders, lumens) are refined with ilastik. Tubules are classified using IF channels and specific markers (LTL, uromodulin, AQP2). Finally, protein and lipid concentrations are measured within the cytoplasm, providing insights into tissue composition. This pipeline enables robust segmentation and quantification of protein/lipid concentrations across anatomical structures, facilitating downstream tissue analysis.

**Fig. 3:**
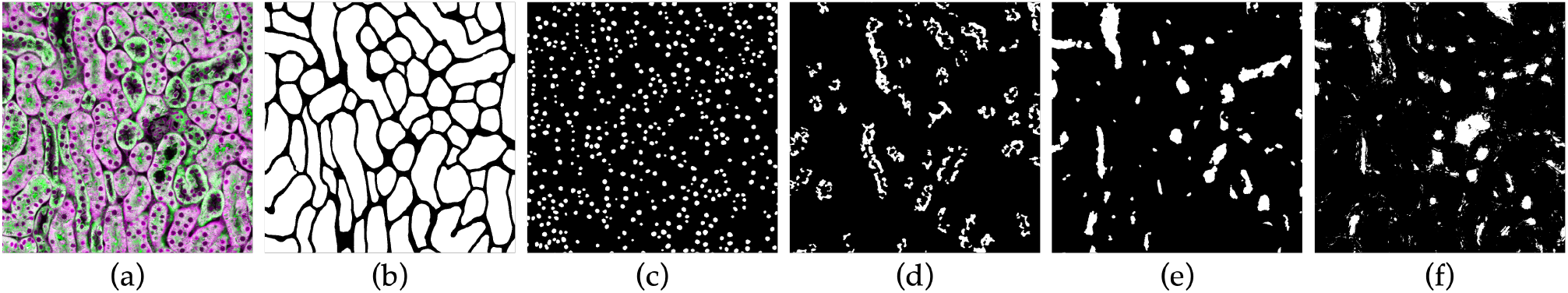
Segmentation of structural and biochemical features in NoRI images. This panel illustrates the full segmentation pipeline applied to NoRI data, enabling downstream quantification of tissue architecture and biochemical content. (a) Raw composite NoRI image-patch showing protein (magenta) and lipid (green) channels. (b) Tubule segmentation mask outlining individual tubules. (c) Nuclei segmentation showing discrete nuclear regions. (d) Brush border segmentation, highlighting apical specializations in proximal tubules. (e) Lumen segmentation, identifying internal tubule spaces. (f) Background and residual lumen segmentation used for masking non-tubular regions.

Initially, raw NoRI images were processed using the Segment Anything Model (SAM) to produce preliminary tubule segmentations. These automated results provided a strong starting point but often included inaccuracies or missed structures. To improve segmentation fidelity, domain experts performed a detailed review and refinement of the initial masks using Fiji (4). This expert intervention corrected systematic errors and resolved ambiguities that the model could not handle reliably.

A final correction pass was then applied to ensure consistency and accuracy across the dataset. This multi-step approach allowed us to efficiently generate robust ground truth annotations suitable for training and validating downstream models.

## 3. Method

This section describes the computational pipeline developed to segment kidney structures and quantify protein and lipid concentrations from NoRI data. **The pipeline is composed of three sequential modules: segmentation, classification, and measurement. Each component was designed to balance biological accuracy, computational efficiency, and ease of adaptation to new imaging contexts**.

First, we applied a hybrid segmentation strategy that leverages the SAM in combination with domain-specific preprocessing and post-processing steps to identify key anatomical structures, including tubules, nuclei, lumens, and brush borders. Next, segmented tubules were classified into anatomical subtypes using IF markers and a machine learning classifier. Finally, protein and lipid concentrations were quantified specifically within the cytoplasmic region of each tubule, excluding nuclei, lumens, and brush borders. This modular framework enables robust, high-throughput analysis of multi-channel tissue images and provides detailed biochemical profiling at single-tubule resolution. All code, datasets, and a visualizer tool are publicly available (see Appendix II for details).

### 3.1. Segmentation

#### 3.1.1. Tubule Segmentation

Tubule segmentation was performed using the SAM (2), a state-of-the-art, vision-transformer-based segmentation model pre-trained on natural image datasets. While SAM offers strong generalization, its direct application to biological tissue images is non-trivial due to several domain-specific challenges: (i) morphological differences between natural and biological images; (ii) incompatibility in data formats (RGB vs. multi-channel 16-bit NoRI); (iii) limited contrast in biological structures; and (iv) the large size of microscopy images, which often exceed GPU memory limits.

To overcome these limitations, we introduced a tailored pipeline consisting of three key adaptations: (1) a domain-specific preprocessing step to generate a third input channel from NoRI data, enhancing structural contrast; (2) a patch-based segmentation strategy with overlapping tiles to handle large images while preserving elongated tubule structures; and (3) a biologically informed post-processing pipeline for refining tubule masks and eliminating segmentation artifacts.

##### Pre-processing

Segmentation relied solely on the two NoRI channels (protein and lipid), ensuring robustness independent of immunofluorescence (IF) markers, which varied across experiments. Each NoRI channel was individually normalized via histogram equalization to improve dynamic range utilization and contrast. **A composite third channel was then created by pixel-wise multiplication of the normalized protein and lipid channels.This operation enhanced boundary definition and improved SAM’s ability to segment tubule structures accurately**.

##### Patch-based Segmentation

Due to the high resolution of NoRI images, full-frame inference with SAM was infeasible. To address this, each image was divided into overlapping patches of 1970 × 2000 pixels with a 500-pixel overlap. This strategy preserved spatial continuity for long tubules crossing tile boundaries. Within each patch, a dense prompt grid (64 points per side) guided SAM’s segmentation, enabling comprehensive coverage of the tubule structures. Redundant masks were suppressed using a non-maximum suppression threshold of 0.9, reducing false positives and improving instance separation.

##### Post-processing

Following initial segmentation, biologically motivated post-processing steps were applied to refine the tubule masks. First, masks occupying more than 30% of a patch or exhibiting low average intensity were discarded as likely non-tubular regions. Valid masks were then merged across patches using logical operations to reconstruct complete tubules. To prevent merged tubules due to close apposition, each tubule mask was eroded using a circular structuring element (radius: 7 pixels). This improved separation between adjacent tubules without distorting their shapes.

Finally, the refined masks were assembled into full-resolution segmentation maps using the 500-pixel overlap to ensure seamless continuity.

##### Expert Validation

To ensure accuracy, all automated segmentations were manually reviewed by a domain expert. Merged tubules were separated, and misidentified structures such as blood vessels or glomeruli were excluded. **This validation step required only 2–5 minutes per image, significantly less than the** ~ **8 hours needed for manual segmentation, making it practical for large-scale datasets**.

#### 3.1.2. Nuclei Segmentation

To segment nuclei, we employed *ilastik*’s machine learning-based pixel classification workflow, using the two processed NoRI channels as input features (5). The classifier was trained on a representative subset of images to distinguish nuclei from background structures, leveraging intensity, texture, and shape features. Once trained, it was applied across the full dataset to generate a binary mask identifying nuclear regions with high confidence.

To eliminate false positives and improve mask quality, a morphological opening was performed using a circular structuring element with a 5-pixel radius. This post-processing step removed small, misclassified objects while preserving the compact, rounded morphology characteristic of nuclei.

#### 3.1.3. Lumen Segmentation

Lumen segmentation also relied on *ilastik*’s pixel classifier, trained to distinguish between luminal and non-luminal regions based on NoRI image features. Due to morphological similarities between lumens and other non-tubular structures (e.g., blood vessels, glomeruli), the classifier was additionally trained to identify these confounding classes as background. This broader training strategy enabled more accurate delineation of tubular lumens.

Following segmentation, we retained only those luminal regions located within the interior of segmented tubules, designating them as true lumen. The remaining areas were designated as background and later used to exclude non-tubular objects from the SAM-generated masks. This refinement step was critical to resolving structural ambiguities introduced by morphologically similar entities in kidney tissue.

#### 3.1.4. Brush Border Segmentation

Brush borders, located on the apical side of proximal tubules, present a segmentation challenge due to their fine structure and variable staining. We used the LTL (Lotus *tetragonolobus* lectin) immunofluorescence channel, which specifically labels brush borders, as input for segmentation. To enhance contrast and correct for uneven illumination, a morphological tophat filter was first applied.

Subsequently, we applied Otsu’s thresholding method (6) to segment high-intensity brush border regions. A final morphological opening with a circular structuring element removed isolated bright pixels and preserved continuous brush border features. This multi-step approach enabled reliable detection of brush borders despite variability in staining intensity and structural morphology.

### 3.2. Classification

In addition to segmentation, classifying the tubule types was essential for investigating subtype-specific differences in protein and lipid concentrations. We combined the tubule masks from the previous step with IF channel data to assign each tubule to a molecular subtype. Specifically, we utilized three markers: LTL (Lotus *Tetragonolobus* Lectin), UMOD (Uromodulin), and AQP2 (Aquaporin 2), which label proximal tubules, thick ascending limbs, and collecting ducts, respectively.

However, due to variability in staining intensity and background fluorescence, directly thresholding marker intensities was unreliable. Furthermore, not all tubules could be confidently assigned to one of the three known types, necessitating a fourth category for unclassified tubules.

To address these challenges, we employed *ilastik*’s object classifier. The model was manually trained on a subset of labeled images, using texture and intensity features to assign tubule classes. Once trained, the classifier was applied to the remaining dataset, excluding ischemic samples.

For ischemic kidneys—where the LTL marker was replaced by a damage-specific marker—the classification task was simplified to binary categorization: ischemia-positive versus negative tubules. Due to lower background fluorescence in these images, a custom filtering-based analysis pipeline was sufficient to assign tubule class labels for downstream analysis.

### 3.3. Measurement

Quantification of protein and lipid concentrations was performed by measuring pixel intensities within the cytoplasm of each segmented tubule. To isolate the cytoplasmic region, masks corresponding to lumens, brush borders, and nuclei were subtracted from the tubule mask, ensuring that only the cytoplasmic compartment was analyzed.

For each resulting cytoplasm mask ℳ, we extracted pixel intensities from the NoRI protein and lipid channels, denoted *I*_(protein)_ and *I*_(lipid)_, respectively. These values were converted to concentrations using established calibration factors: 1.3643 for protein (based on bovine serum albumin, BSA) and 1.0101 for lipid (based on DOPC). The total concentrations, expressed in mg/ml, were computed as:

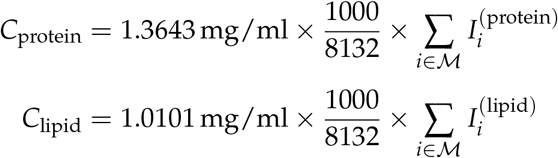

Here, the factor 1000/8132 accounts for unit normalization based on the imaging system’s calibration. These calculations resulted per-tubule protein and lipid concentrations, facilitating a detailed biochemical assessment across tubule subtypes and experimental conditions. This allows detailed molecular profiling of individual tubules, supporting quantitative comparisons across conditions.

## 4. Results

### Across 100 images, we identified 37,622 tubules and 213,645 nuclei

Among the segmented tubules, 80.59% contained a lumen, and 30.70% exhibited brush borders. These statistics, along with subtype-specific protein and lipid measurements, highlight the pipeline’s ability to extract detailed morphological and molecular information across diverse kidney samples. Further biological analyses across experimental groups are presented in the accompanying biological study (3).

### Quantitative analysis of segmented tubules revealed distinct biochemical profiles across renal tubule subtypes

As shown in Fig. **4**, tubules positive for LTL and Uromodulin exhibited the highest mean protein and lipid concentrations, reflecting their roles in reabsorption and active transport. AQP2-positive collecting ducts displayed markedly lower concentrations, consistent with their reduced metabolic activity. Unlabeled tubules showed broader variability, potentially representing a heterogeneous mixture of tubule types or classification uncertainty. **These findings demonstrate that the pipeline enables subtype-specific biochemical characterization at single-tubule resolution. These findings confirm the utility of cytoplasm-resolved biochemical profiling for differen-tiating renal tubule subtypes**.

**Fig. 4:**
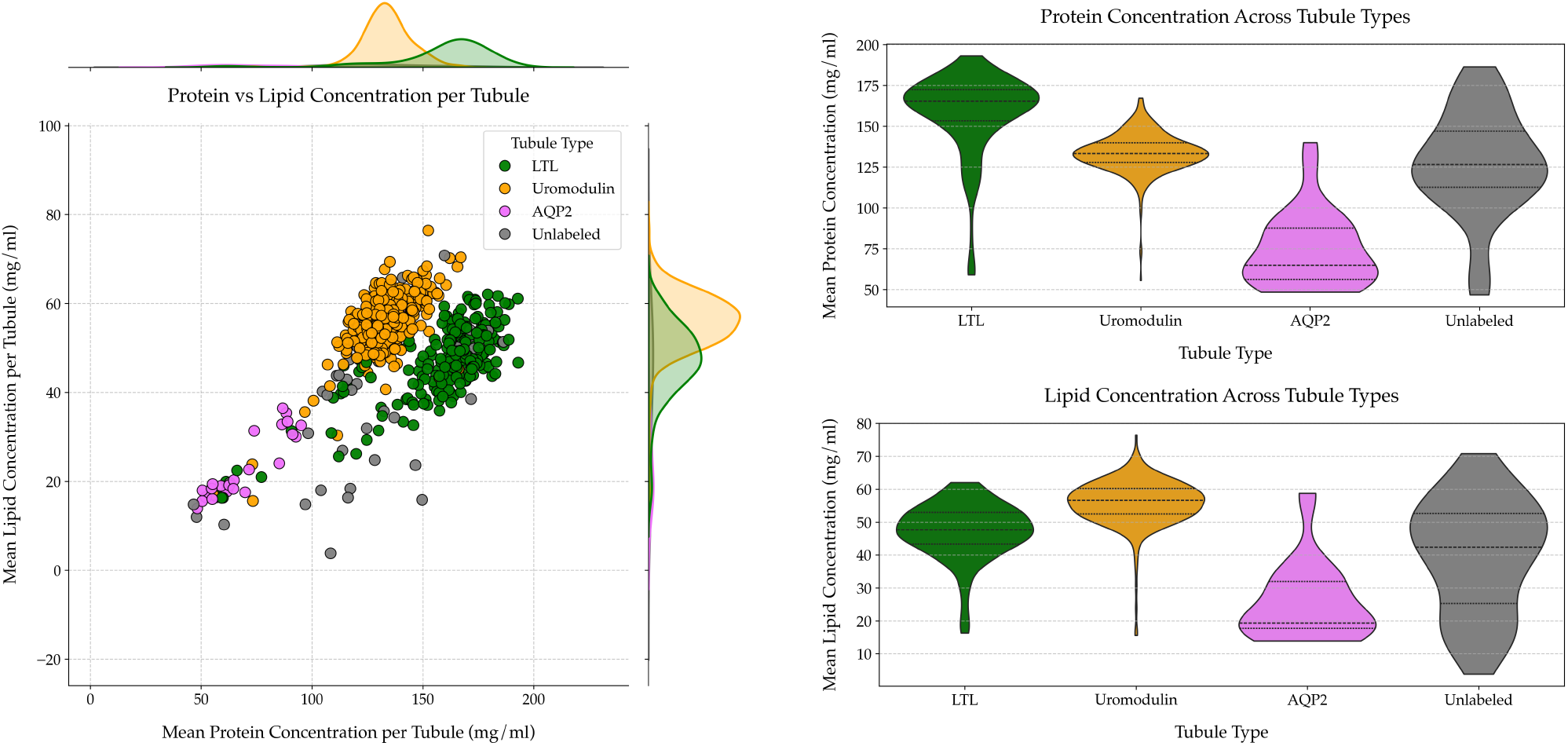
Quantitative biochemical profiling of renal tubule subtypes using NoRI. (Left) Joint distribution of mean protein and lipid concentrations per tubule reveals distinct clusters by molecular marker, indicating characteristic biochemical signatures for LTL-, Uromodulin-, and AQP2-positive tubules. (Right, top) Protein concentrations are highest in LTL and Uromodulin tubules, and lowest in AQP2-positive collecting ducts. (Right, bottom) Lipid concentrations are similarly elevated in Uromodulin-positive tubules and reduced in AQP2 tubules. Unlabeled tubules exhibit broad variability across both modalities. These results highlight NoRI’s utility for label-free biochemical classification of renal tubules.

**Fig. 5:**
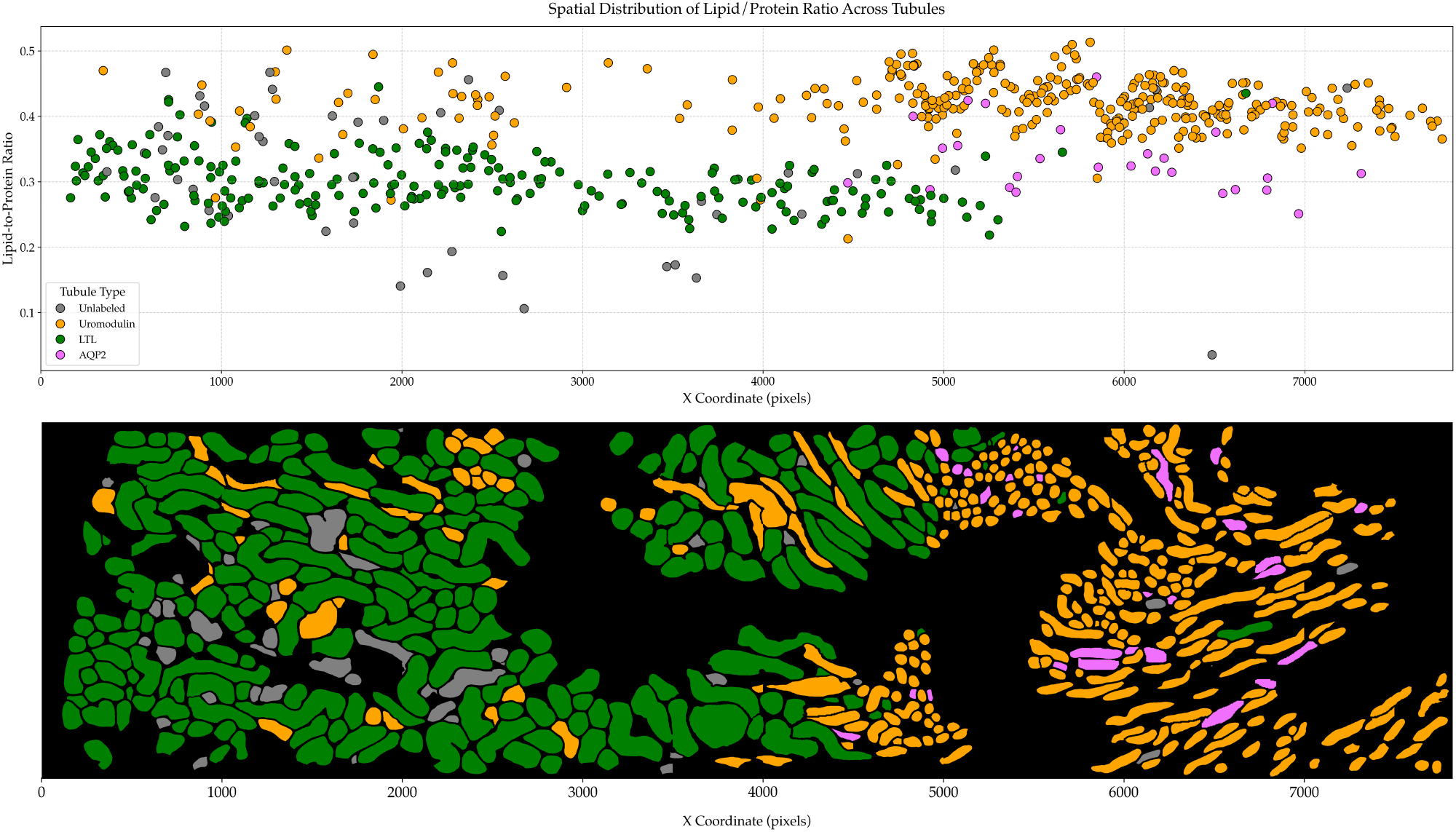
Spatial distribution of lipid-to-protein ratios reveals regional variation in tubule biochemistry. (Top) Scatter plot of the lipid-to-protein ratio for each segmented tubule, plotted against its X-coordinate in the tissue section. Uromodulin-positive tubules exhibit consistently higher ratios compared to LTL- and AQP2-positive tubules, with a sharp spatial transition near X = 5000 pixels. (Bottom) Corresponding segmented tissue map showing the spatial organization of tubule types. Uromodulin-positive tubules (orange) dominate the right region, correlating with elevated lipid-to-protein ratios, while LTL-positive tubules (green) are more abundant in the left region. AQP2-positive collecting ducts (magenta) are scattered and maintain lower ratios. This demonstrates NoRI’s capability to reveal tissue-level biochemical heterogeneity and its anatomical context.

From a computational perspective, **the full pipeline achieved an F1 score of 0.9226 for tubule segmentation, with strong agreement to manual ground truth**. The use of a custom contrast-enhanced third channel, in combination with SAM and ilastik-based refinement, contributed to high segmentation accuracy and robustness. The tubule segmentation using SAM was the most expensive step in the pipeline which took about 20 seconds per image patch of 2000 × 2000 on a consumer grade GPU. Running the full pipeline takes between 1 and 5 minutes on the same hardware, depending on the image size. **This shows that the pipeline can be run on consumer grade computer or laptop with almost real-time analysis results**.

The segmentation pipeline developed in this study served as the analytical foundation for the biological investigation presented in Trim et al. (3). By enabling precise delineation of kidney tubules and substructures (nuclei, lumen, brush border, and cytoplasm) from multi-channel NoRI images, the pipeline allowed cytoplasm-resolved quantification of protein and lipid concentrations at single-tubule resolution. These quantitative maps revealed distinct biochemical signatures between renal tubule subtypes, with proximal (LTL+) tubules showing high protein and moderate lipid densities, distal (Uromodulin+) tubules exhibiting lower protein but higher lipid content, and collecting ducts (AQP2+) displaying the lowest values across both modalities. Importantly, the segmentation accuracy and reproducibility of this framework enabled robust statistical comparisons across sex, anatomical regions, and disease states, forming the quantitative basis for discovering several novel physiological and pathological patterns.

When integrated with machine learning pipelines (e.g., ResNet50 and YOLO-based models), the segmentation outputs allowed large-scale, automated feature extraction from NoRI datasets, revealing sex-specific and injury-dependent biochemical trends with high confidence. For instance, analyses of ischemia–reperfusion injury showed dynamic remodeling of protein and lipid distributions, identifying day 2 post-injury as a critical inflection point with sharp reductions in cytoplasmic lipid concentration and brush border disruption—features that later recovered during tissue repair. Similarly, sex-differentiated protein and lipid distributions were detectable from regions as small as 130 µm^2^, underscoring the precision and consistency of the computational pipeline. These findings highlight how the segmentation framework not only accelerated quantitative histopathology but also enabled the biological interpretation of NoRI data, bridging computational imaging and mechanistic insight into kidney physiology and pathology.

## 5. Discussion

We present a biologically consistent and computationally efficient pipeline that adapts general-purpose vision models for the quantitative analysis of NoRI kidney images. By integrating state-of-the-art segmentation tools with domain-specific enhancements, this framework enables automated segmentation and quantification of kidney structures with high accuracy and scalability. The resulting pipeline is fully modular, supports large-scale analysis, and facilitates detailed biochemical profiling of individual renal tubules across diverse experimental conditions.

A principal contribution of this framework is its capacity for cytoplasm-resolved quantification of protein and lipid concentrations within individual tubules. By explicitly removing nuclei, lumens, and brush borders, the method isolates cytoplasmic regions and generates spatially resolved molecular profiles. These measurements facilitate high-resolution analysis of tissue heterogeneity and enable biologically meaningful applications, such as assessing tubule-specific injury responses or regional biochemical variation within the kidney.

A defining feature of the pipeline is its modular design, which supports independent modification of the segmentation, classification, and quantification components. This flexibility allows the framework to be adapted to other tissue types imaged using NoRI with minimal reconfiguration. Notably, the segmentation module relies exclusively on the two Raman channels, avoiding dependence on immunofluorescence (IF) markers, which are often variable across experiments. This choice ensures robustness across imaging conditions and improves generalizability across datasets.

The pipeline is also optimized for computational performance. Image tiling with overlapping patches allows for the use of transformer-based models such as SAM while avoiding memory bottlenecks. Subsequent post-processing and ilastik-based refinement improve segmentation quality with minimal computational overhead. On a standard GPU-equipped workstation, the full pipeline processes high-resolution images in under a minute. On CPU-only systems, the pipeline remains functional, albeit with increased processing times, offering adaptability across computational environments.

The system achieves high segmentation accuracy, with an F1 score of 0.9226, and demonstrates strong concordance with expert-curated ground truth. Crucially, biological measurements derived from automated segmentations are indistinguishable from those obtained using manual annotations, both in tubule counts and in quantification of protein and lipid concentrations. This validates the pipeline’s capacity to deliver biologically faithful results in a fully unsupervised manner.

Each component of the pipeline was rigorously evaluated through controlled experiments (Appendix I). Preprocessing strategies, third-channel construction methods, encoder configurations, and post-processing steps were systematically compared. These ablations demonstrated that each design choice—such as the use of a contrast-enhancing third channel or ilastik-based filtering—contributes measurably to the observed accuracy and consistency of results.

Reproducibility was a central design objective throughout the pipeline (7). All computational steps are deterministic and use open-source tools, ensuring transparency, accessibility, and ease of integration into broader bioimage analysis workflows. Outputs are fully compatible with standard platforms, facilitating visualization and downstream analysis.

In summary, this work introduces a robust, generalizable, and reproducible computational framework for the quantitative analysis of NoRI kidney images. It illustrates how domain-specific adaptations of generalist vision models, coupled with principled algorithmic choices, can enable biologically meaningful and scalable tissue analysis. The pipeline’s ability to deliver tubule-resolved biochemical measurements with high fidelity and speed makes it a valuable tool for both research and translational applications.

## 6. Conclusion

Developing this pipeline highlighted both the challenges and opportunities of working with large, multi-channel NoRI datasets. Instead of building new models from scratch which is often data-intensive, requires extensive training, and depends on large annotated datasets, we found that combining domain knowledge with careful design choices allowed us to use existing tools more effectively. In particular, adapting a foundation model like SAM with targeted preprocessing and post-processing steps resulted in accurate and scalable segmentation suited to our specific imaging modality.

This approach shows that with thoughtful integration, general-purpose models can be applied to specialized biological data without the need for custom training. It also reinforces the importance of designing workflows that are modular, reproducible, and easy to adapt as experimental conditions or tissue types change.

Looking ahead, the same strategy could be extended to other imaging modalities or analysis goals. As datasets continue to grow in size and complexity, approaches like this may help make quantitative, label-free analysis more practical and broadly usable across biomedical research.

## Appendix I – Experiments

In this section, we systematically explore the impact of various components in the segmentation pipeline on the accuracy of kidney tubule segmentation, utilizing a detailed and structured approach. We conducted experiments that cover three critical areas: the creation of the third channel, the selection of image encoders within SAM, and the necessity of specific pre-processing and post-processing steps.

### Experiment 1: Preprocessing Ablations - Third-Channel Construction

The Segment Anything Model (SAM) requires a three-channel image input. Since NoRI imaging provides only two channels—protein and lipid—we investigated multiple strategies for constructing a third channel tailored to biological tissue segmentation. The aim was to improve contrast and boundary detection in SAM’s input space while preserving biological fidelity.

We evaluated eight different third-channel configurations:

- **[Protein, Protein, Protein]**: Monochromatic triplet using protein channel only.
- **[Lipid, Lipid, Lipid]**: Monochromatic triplet using lipid channel only.
- **[Protein, Lipid, Protein]**: Interleaved configuration favoring protein signal.
- **[Protein, Lipid, Lipid]**: Interleaved configuration favoring lipid signal.
- **[Protein, Lipid, Endomucin]**: Using an immunofluorescence channel as the third input.
- **[Protein, Lipid, Zeros]**: All-zero third channel baseline.
- **[Protein, Lipid, Average]**: Mean of the two channels.
- **[Protein, Lipid, Custom]** (proposed): Pixel-wise multiplication of protein and lipid channels, with histogram equalization to enhance boundary contrast.

As shown in Figure **6**, the choice of third channel has a marked impact on segmentation quality. The custom method consistently outperformed all other variants, achieving the highest mean F1 score with the lowest variance. This improvement is statistically significant across most pairwise comparisons (*p <* 0.001). The pixel-wise multiplication strategy accentuated co-localized signal regions, enhancing edge definition critical for tubule boundary segmentation.

**Fig. 6:**
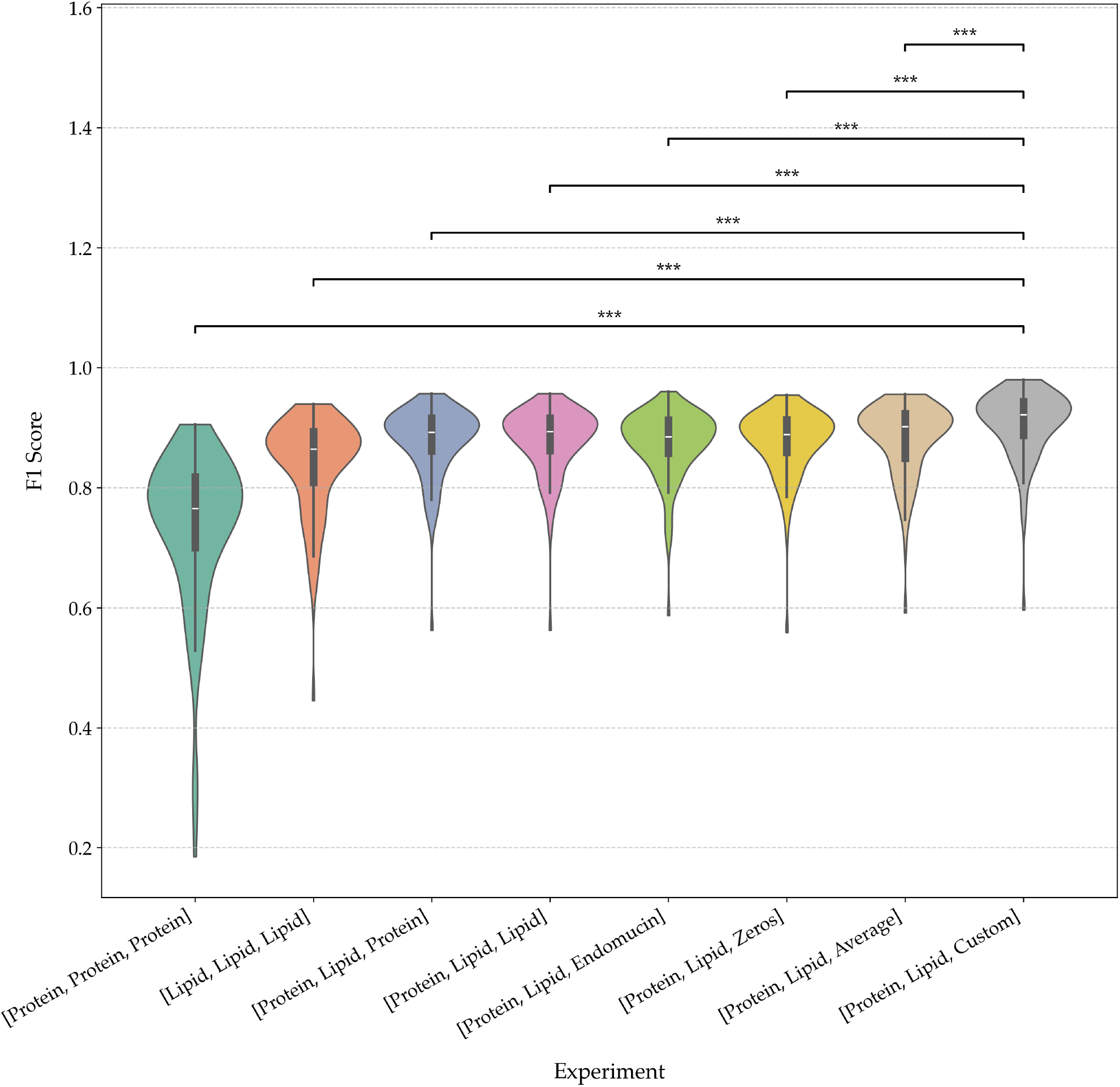
Comparison of segmentation performance across NoRI channel combinations. Violin plots of F1 scores from segmentation models trained on different combinations of NoRI input channels. The baseline models using only protein or only lipid channels underperform compared to those using both. Replacing the third input channel with either synthetic (Zeros), averaged, or domain-specific (Custom) data significantly improves performance, with the best results observed for the [Protein, Lipid, Custom] configuration. Asterisks denote statistically significant differences (*** *p <* 0.001, ** *p <* 0.01), highlighting the impact of channel selection on segmentation quality. This demonstrates that biologically informed channel construction significantly improves segmentation fidelity.

These results emphasize the importance of biologically motivated preprocessing when adapting generalist models to domain-specific imaging modalities. The success of the custom channel supports its use as a default strategy in future NoRI-based segmentation tasks.

### Experiment 2: Evaluation of SAM Image Encoders

The SAM model offers three different image encoders—vit-b, vit-l, and vit-h—that vary significantly in their architectural complexity and computational demands:

- **vit-b:** A base encoder with 91 million parameters.
- **vit-l:** A large encoder with 308 million parameters.
- **vit-h:** A huge encoder with 636 million parameters.

Each of these encoders was evaluated based on their ability to segment kidney tubules accurately and the computational resources required to do so. As shown in Figure **7**, the vit-h encoder consistently delivered the most accurate segmentation, with an F1 score of approximately 0.9264, surpassing the performance of both vit-b and vit-l. While the vit-h encoder is computationally intensive—requiring approximately 27 additional seconds per image on a GPU compared to the smaller encoders—this increase in processing time is justified by the significant improvement in segmentation accuracy. This trade-off is particularly relevant in research and clinical settings where precision is paramount, and computational resources are available. The superior performance of the vit-h encoder can be attributed to its ability to better capture and process the complex features within the images, thus leading to more accurate segmentation.

**Fig. 7:**
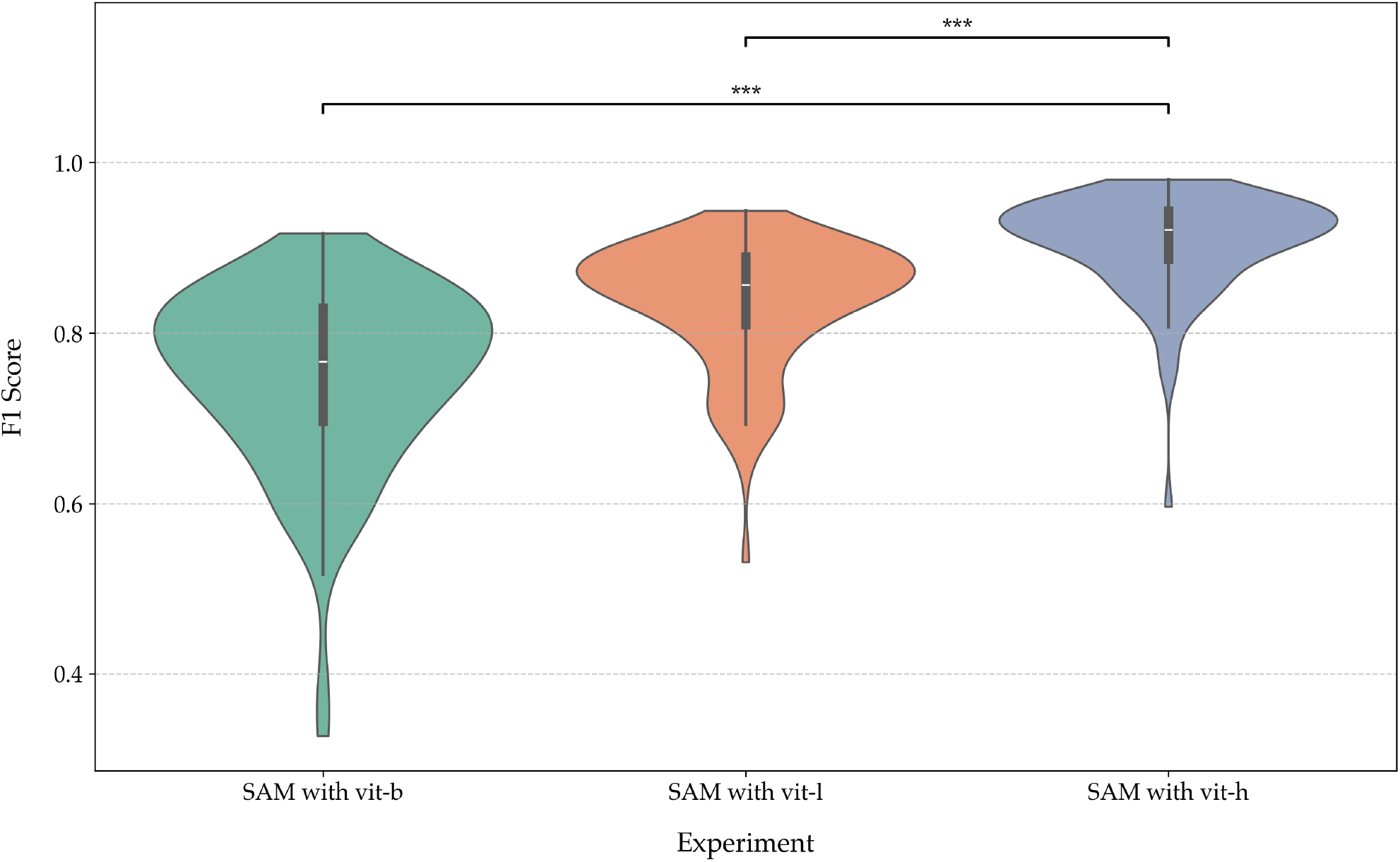
Experimenting with Different Image Encoders in SAM: Violin plot illustrating the F1 score distributions for experiments with different image encoders in SAM. The configurations include vit-b (base), vit-l (large), and vit-h (huge). The plot highlights the variability and central tendencies in performance for each encoder type.

### Experiment 3: Ablation Study on the Segmentation Pipeline

To understand the role of each step in the segmentation pipeline, an ablation study was performed (Figure **8**), where key components were systematically omitted:

**Fig. 8:**
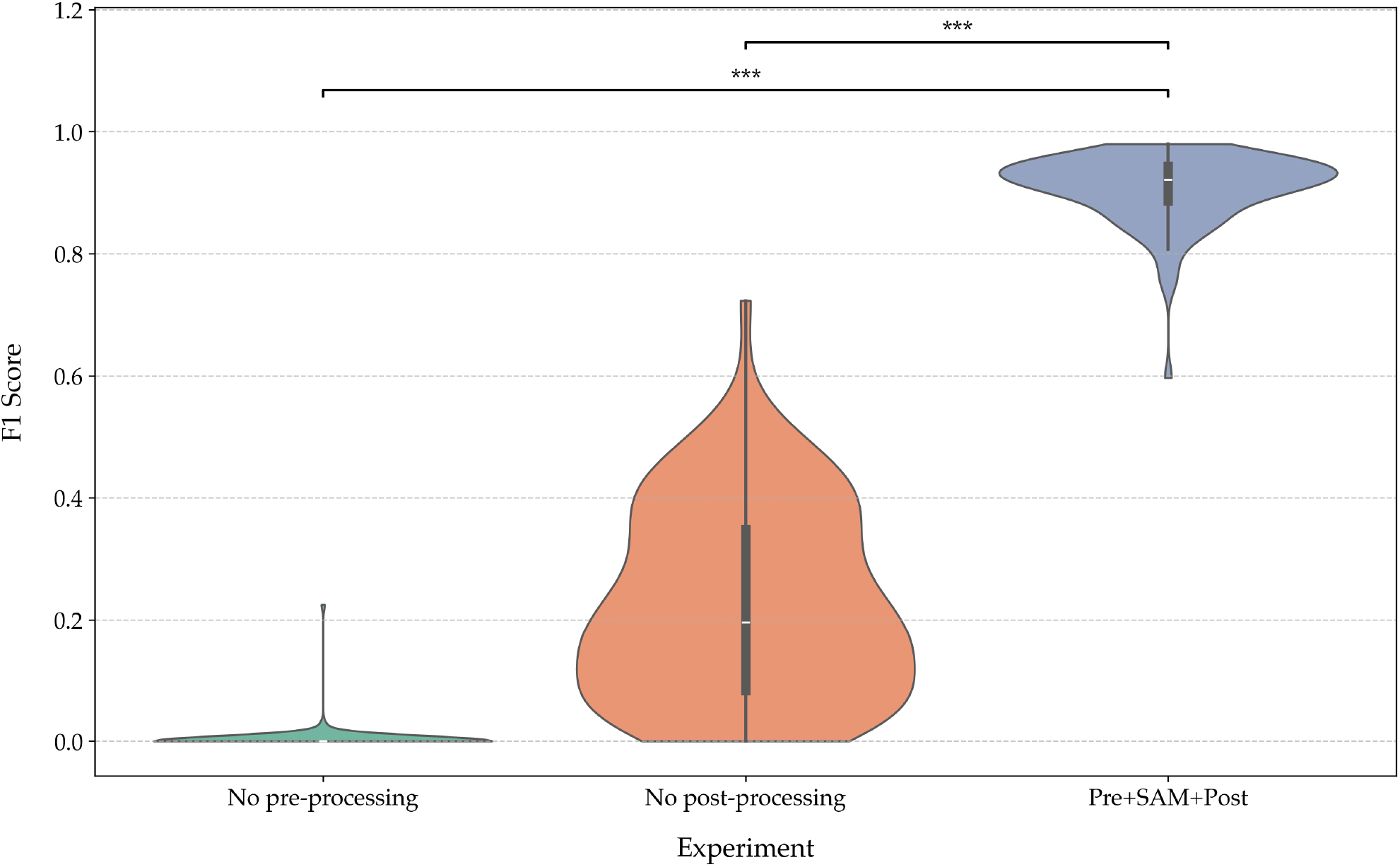
Ablation study evaluating the impact of different processing steps on segmentation performance. Violin plot showing the distribution of F1 scores across experiments with different processing steps. The experiments include no pre-processing, no post-processing, the combination of pre-processing, SAM segmentation, and post-processing, and the addition of ilastik cleaning. This analysis highlights the impact of each step on model performance, demonstrating the importance of a complete pipeline.

**Fig. 9:**
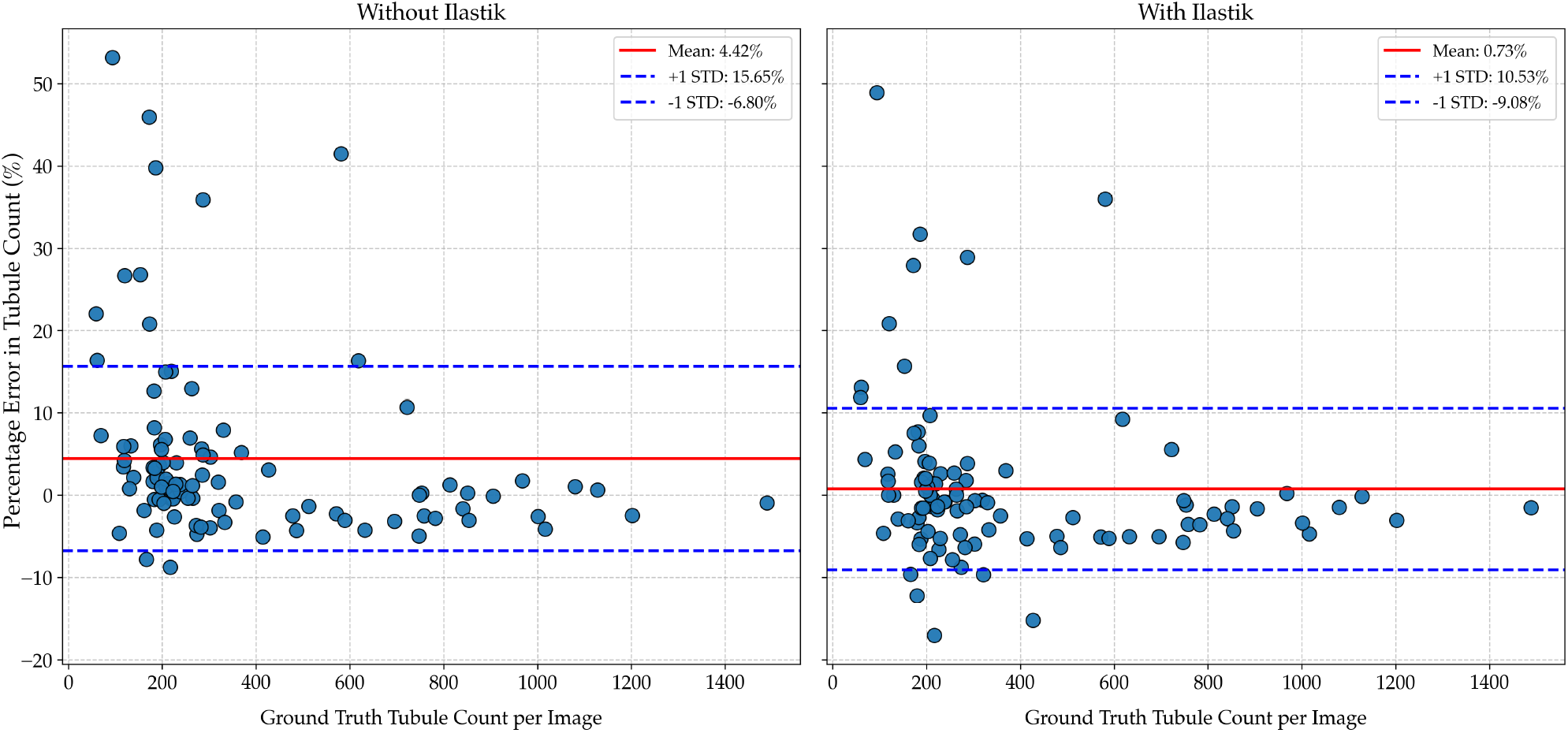
Impact of ilastik filtering on tubule count accuracy. Percentage error in automatic tubule counts compared to ground truth, shown for models without (left) and with (right) ilastik-based filtering. Without ilastik, the mean error is higher (4.42%) with greater variability, particularly in low-count images. Incorporating ilastik reduces both the mean error (0.73%) and standard deviation, improving consistency across the full range of tubule densities. These results demonstrate that ilastik filtering enhances segmentation accuracy, especially in challenging regions with sparse or ambiguous tubules.

- **Preprocessing Omission:** Without preprocessing, the segmentation process failed entirely, indicating that preprocessing is critical for the model’s ability to correctly identify and segment kidney tubules. The preprocessing step, which includes histogram equalization and the creation of the third channel, enhances the contrast and quality of the input images, making the features more discernible for the SAM model.
- **Post-processing Omission:** The omission of post-processing steps led to a significant decrease in segmentation accuracy. The segmented tubules appeared merged and less distinct, highlighting the importance of post-processing in refining the segmentation results. Post-processing steps are crucial for cleaning up the initial segmentation by removing noise and non-tubule elements, thereby ensuring that only the relevant structures are retained.
- **ilastik Refinement:** The inclusion of ilastik for final cleanup significantly improved the segmentation quality, increasing the average F1 score from 0.9212 to 0.9226. While this improvement may seem modest, it is particularly impactful for images that initially had lower accuracy. The use of ilastik introduces a pixel-based classification that further refines the segmentation, reducing variability and enhancing the consistency of the results across the dataset.

These results indicate that protein and lipid concentrations measured from segmentations are statistically robust across variations in the pipeline. These small effect sizes confirm that automation does not introduce meaningful bias, validating the pipeline’s use for biological inference. The ablation study clearly demonstrates that each component of the pipeline is essential and contributes uniquely to the overall accuracy of the segmentation task.

### Experiment 4: Comparing biological outcome - ground truth vs fully automatic

To assess the impact of segmentation accuracy on downstream biological measurements, we compared protein and lipid concentrations obtained from three segmentation approaches: manual ground truth, fully automated segmentation using SAM, and the automated pipeline with ilastik refinement. This comparison was performed to determine whether deviations in segmentation accuracy influence the resulting biochemical measurements.

The results indicate that protein and lipid concentrations measured across these methods are highly consistent. The effect size between manual and fully automated segmentation was small (Cohen’s d = –0.053), suggesting that the differences introduced by automation do not substantially alter the biochemical readouts. With the addition of ilastik-based refinement, the effect size compared to the manual segmentation further decreased (Cohen’s d = 0.008). The difference between the SAM-only and SAM+ilastik methods was similarly modest (Cohen’s d = –0.046).

These findings indicate that although there are small differences in segmentation masks as reflected by F1 score, the overall conclusions regarding protein and lipid distributions remain unchanged. The automated pipeline can therefore be used with confidence for large-scale analysis where manual annotation or correction is not practical.

The results from these experiments collectively provide a comprehensive understanding of the factors influencing kidney tubule segmentation accuracy using the SAM model. The creation of a third channel through a custom method involving multiplication and preprocessing proved to be the most effective approach, highlighting the importance of enhancing contrast and combining multiple data sources to improve segmentation quality.

The evaluation of different SAM encoders revealed that the vit-h encoder, despite its higher computational cost, offers the best performance in terms of accuracy, making it the preferred choice when resources allow. The ablation study underscored the necessity of both preprocessing and post-processing steps, with each playing a crucial role in achieving high segmentation accuracy. The refinement introduced by ilastik, although seemingly minor, significantly enhances the consistency of the results, particularly for more challenging images.

Overall, this study presents a robust and reliable segmentation pipeline that, when fully implemented, achieves an F1 score of 0.9226, closely aligning with the ground truth. These findings not only validate the chosen methodology but also provide a solid foundation for further research and application in biomedical image analysis, particularly in the accurate segmentation of kidney tubules. The insights gained from this study can inform future efforts to optimize segmentation pipelines and improve the accuracy of automated image analysis in complex biological datasets.

## Appendix II – Code, Data and Visualizer

To support reproducibility and encourage further exploration, we have made all relevant resources publicly available. The full codebase used for our analysis can be accessed on GitHub, along with a dedicated repository for the NoRI visualizer that enables detailed exploration of individual tubules. Additionally, the dataset used in our experiments has been uploaded to Zenodo.

- **Pipeline GitHub:** https://github.com/HMS-IAC/nori
- **Visualizer GitHub:** https://github.com/HMS-IAC/NoRI-Visualizer
- **Dataset Zenodo:** https://doi.org/10.5281/zenodo.17329675
- **Project website:** https://hms-iac.github.io/NoRI/

## Acknowledgments

We gratefully acknowledge Federico Gasparoli for his contributions to the NoRI visualizer. His assistance in developing and refining the visual interface was critical to the clarity and effectiveness of our results. We also thank Kseniia Petrova for her essential support in data collection and for her guidance on data-related tasks.

## References

[1] S. Oh, C. Lee, W. Yang, A. Li, A. Mukherjee, M. Basan, C. Ran, W. Yin, C. J. Tabin, D. Fu, et al., “Protein and lipid mass concentration measurement in tissues by stimulated raman scattering microscopy,” Proceedings of the National Academy of Sciences, vol. 119, no. 17, p. e2117938119, 2022.

[2] A. Kirillov, E. Mintun, N. Ravi, H. Mao, C. Rolland, L. Gustafson, T. Xiao, S. Whitehead, A. C. Berg, W.-Y. Lo, et al., “Segment anything,” in Proceedings of the IEEE/CVF International Conference on Computer Vision, pp. 4015–4026, 2023.

[3] W. V. Trim, S. Oh, M. Diakova, K. Petrova, T. Ichimura, A. Takakura, R. Karmakar, S. F. F. Norrelykke, L. Peshkin, J. V. Bonventre, and M. W. Kirschner, “Normalized raman imaging for studies of tissue physiology of the kidney,” bioRxiv, 2025.

[4] J. Schindelin, I. Arganda-Carreras, E. Frise, V. Kaynig, M. Longair, T. Pietzsch, S. Preibisch, C. Rueden, S. Saalfeld, B. Schmid, et al., “Fiji: an open-source platform for biological-image analysis,” Nature methods, vol. 9, no. 7, pp. 676–682, 2012.

[5] S. Berg, D. Kutra, T. Kroeger, C. N. Straehle, B. X. Kausler, C. Haubold, M. Schiegg, J. Ales, T. Beier, M. Rudy, et al., “Ilastik: interactive machine learning for (bio) image analysis,” Nature methods, vol. 16, no. 12, pp. 1226–1232, 2019.

[6] N. Otsu et al., “A threshold selection method from gray-level histograms,” Automatica, vol. 11, no. 285-296, pp. 23–27, 1975.

[7] K. Miura and S. F. Nørrelykke, “Reproducible image handling and analysis,” The EMBO journal, vol. 40, no. 3, p. e105889, 2021.

